# Trisk 95 as a Novel Skin Mirror for Normal and Diabetic Systemic Glucose Level

**DOI:** 10.1101/688887

**Authors:** Nsrein Ali, Hamid Reza Rezvani, Diana Motei, Sufyan Suleman, Walid Mahfouf, Isabelle Marty, Veli-Pekka Ronkainen, Seppo J. Vainio

## Abstract

Coping with diabetes requires frequent and even today mostly invasive blood glucose-based monitoring. Partly due to this invasive nature and the associated reduced skin wound healing and increased risk of infection, non-invasive glucose monitoring technologies would represent considerable progress. Edited keratinocytes may enable such a function.

To address this hypothesis, we conducted a proteomic screen in the skin by making use of the experimental *in vivo* mouse model of type I diabetes alongside controls. We identified Trisk 95 as the only protein whose expression is induced in response to high blood glucose. A luciferase reporter assay demonstrated that induction of Trisk 95 expression occurs not only at the protein level but also transcriptionally. This induction was associated with a marked elevation in the Fluo-4 signal, suggesting a role for intracellular calcium changes in the signalling cascade. Strikingly, these changes lead concurrently to fragmentation of the mitochondria. As judged from the knockout findings, both the calcium flux and the mitochondrial phenotype were dependent on Trisk 95 function, since the phenotypes in question were abolished.

The data demonstrate that the skin represents an organ that reacts robustly and thus mirrors changes in systemic blood glucose levels. The findings are also consistent with a channelling model of Trisk 95 that serves as an insulin-independent but glucose-responsive biomarker taking part in releasing calcium from the cellular stores in the skin. The skin cells may thus provide a novel mean for glucose monitoring when analysing changes in labelled Trisk 95 and calcium. By that, this study is the first proof of the concept of our registered patent (No. PCT FI2016/050917), which proposes the use of cells as biosensors for developing personalized health-monitoring devices.

## Introduction

In view of its dynamic behaviour in the presence of multiples stress, the human skin is widely used to test cellular and molecular responses to specific treatments. Since the skin is one of main organs that are predisposed towards developing damage in diabetic patients, so that between 1/3 and nearly all diabetic patients experience cutaneous complications (1–4), the developing of a non-invasive device for sensing glucose by means of the skin has become a focus of interest for scientists and commercial companies worldwide (5–8). In fact, the currently most widely used devices require skin penetration as far as the interstitial fluid, and are thereby invasive, although a new generation of devices based on the measurement of various metabolites in sweat such as sodium, lactate, potassium and glucose has been proposed recently (9–12). One major drawback with such devices, however, is the amount of sweat required, resulting in secondary effects such as skin irritation. In view of all the devices available on the market and the unknown biological behaviour of skin in response to high glucose, we propose a new paradigm that operates by converting the skin cells to serve as biosensors for monitoring blood glucose.

Keratinocytes are the main cell type (≈90 %) constituting the epidermis (13, 14), and with respect to their key roles in wound healing (15), they have been extensively studied, notably in cases of diabetes (16–18). Reports have demonstrated that impaired keratinocyte function can result in delayed wound healing, and that multiple physiological processes in keratinocytes, such as proliferation (19), migration (20), apoptosis (21) and differentiation (22), may be affected by a hyperglycaemic condition. Moreover, the skin barrier dysfunction (23, 24) and increased inflammation (25–27) brought about by a hyperglycaemic environment will prevent the keratinocytes from healing, probably leading to continuously infected severe skin lesions (28, 29). In view of the literature, keratinocytes could be used as indicators of the status of normal and diabetic skin.

We set out here to investigate whether blood glucose concentration could be monitored by means of keratinocytes and whether a specific signalling pathway is activated by this sensing. Our results show that Trisk 95 expression is increased in the skin and its primary keratinocytes following an increase in blood glucose concentration. In turn, upregulation of Trisk 95 is associated with an increase in the intracellular calcium level, which then triggers a particular modification in the morphology of the mitochondrial network. Our data demonstrate that a specific signalling pathway is activated in the skin and its primary cells in response to a high glucose concentration. These results lead us to propose that keratinocytes could serve as promising biosensors for developing a new generation of blood glucose monitoring devices, especially in diabetic patients.

## Results

### Skin glucose levels are rapidly influenced by glucose concentrations in the blood

Little is known about the responses of epidermal cells to increased blood glucose concentrations or whether a particular signalling network is activated in these cells. To test this hypothesis, we first injected glucose into healthy and type I diabetic mice, the latter having been prepared as previously described (30). Briefly, C57BL/6 mice were divided into two groups: (healthy) controls, which were injected intraperitoneally with citrate buffer (pH 4.5), and type I diabetic mice, which received one single dose of 150 mg/kg streptozotocin (STZ). Blood glucose was monitored every 48 hours and the mice were considered diabetic when its level was ≥ 15 mmol/l (data not shown). The first step was to select a suitable end-point at which the blood glucose level in the healthy mice reaches that found in the diabetic mice (≥ 15 mmol/l), as this can be used for collecting the skin samples. For this purpose, we performed a glucose tolerance test (GTT) on the healthy mice. After fasting for 12 hours, the mice were divided into two groups: a control group in which PBS was injected intraperitoneally, and a treatment group that received 2gr/kg D-glucose. Serum was collected from both groups (15 animals/group) at time-points 0, 5, 15, 30 and 45 min and blood glucose levels were monitored. The results showed a significant increase in blood glucose at 5 min post-injection in the glucose-injected mice (15.20 mmol/l±1.36) (Fig. 1a, full circle) as compared with the placebo-treated mice (6.27 mmol/l±0.31) (Fig. 1a, empty circles). The glucose concentration remained at this level up to the 45 min time-point and then started to drop (Fig. 1a).

**Fig. 1.**
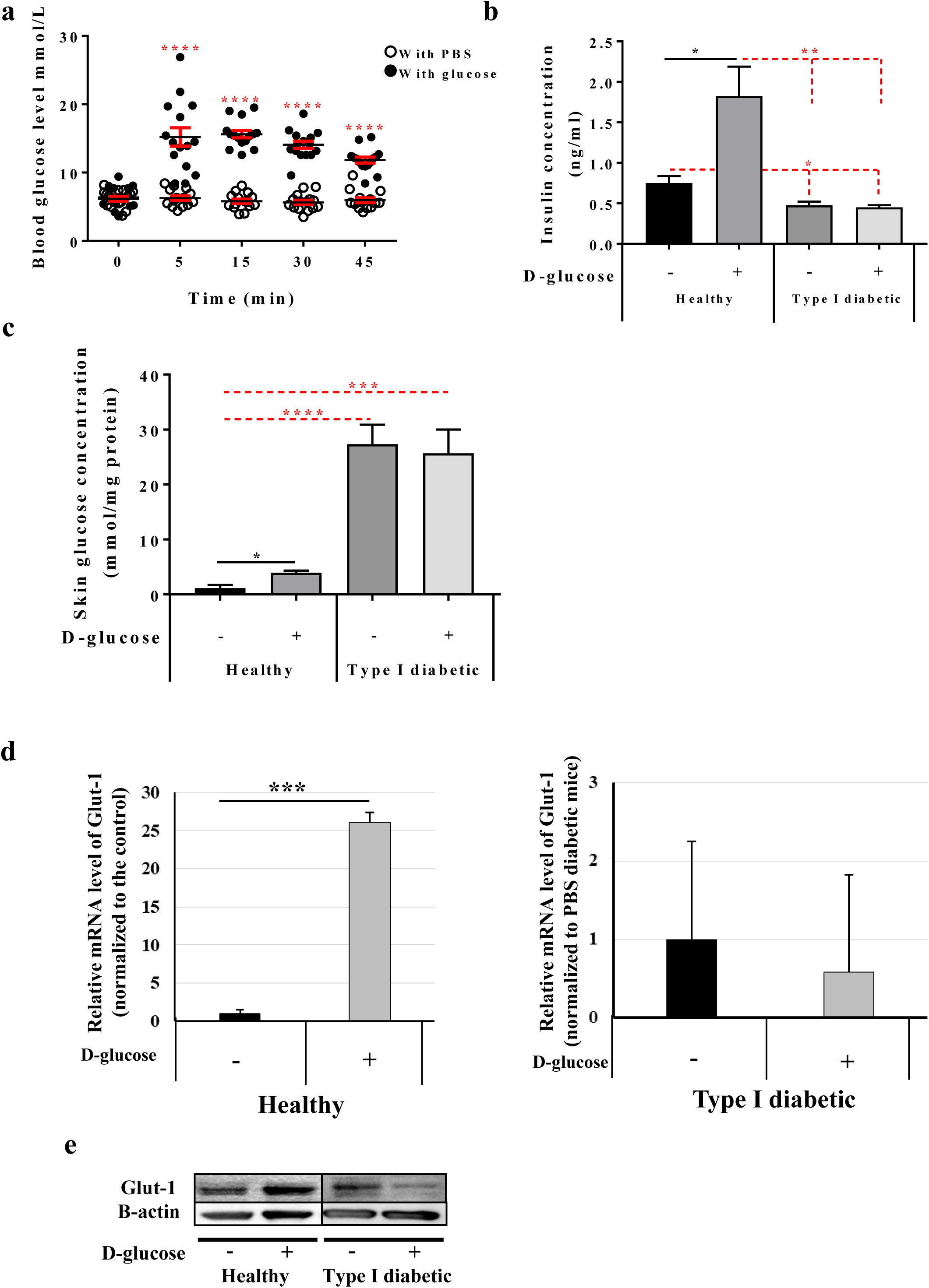
Skin rapidly senses any increase in blood glucose. **a** GTT assay using C57BL/6 mice. The mice were divided into two groups: the control group (empty circle) was injected IP with PBS, and the treated group (full circle) with 2gr/kg D-glucose diluted in PBS. Blood glucose levels were monitored at the given time points (n=15 mice/group). **b** Insulin concentrations in healthy and type I diabetic mice. Insulin levels were measured using the Insulin Rodent Chemiluminescence ELISA technique (n=6 mice/group). **c** Skin glucose levels *in vivo*. The GTT assay was performed and skin biopsies were collected from the four groups (n=5 mice/group). Glucose levels were measured using the YSI 2950 Biochemistry Analyzer (YSI Life Sciences) and normalized to the total concentration of proteins (mmol/mg protein). **d** *Glut-1* mRNA levels in the skin of both healthy and type I diabetic mice was quantified by qRT-PCR. The results are shown as averages after normalization to the controls ± SD (n=6/group). **e** Glut-1 protein expression was determined by western blotting and b-actin was used as a loading control. Two-way ANOVA analysis was performed in a using GraphPad Prism software, **** *P* <0.0001 (a). The two-tailed Student’s t-test was used in b (* *P* <0.02, ** *P* <0.001, ns. not significant 0.25), in c (* *P* <0.02, **** *P* <0.0001 and *** *P* <0.0005, ns. not significant 0.77) and in d (* *P* <0.02 and *** *P* <0.0001).

Blood insulin measurements showed a significant increase upon the injection of glucose in the healthy mice (1.8 ng/ml±0.76 versus 0.83 ng/ml±0.076 in the placebo-injected mice), whereas the blood insulin concentration in the type I diabetic mice did not differ between those receiving a placebo or a glucose injection (0.25 ng/ml±0.038; and 0.20 ng/ml ±0.026 respectively) (Fig. 1b). Although the blood insulin concentration in both the placebo and glucose-injected type I diabetic mice was significantly lower than in their healthy counterparts (Fig. 1b), a certain background level of insulin was detected in both, suggesting that the STZ treatment did not destroy all the β-cells.

To determine whether the skin glucose level changes upon an increase in blood glucose concentration, skin and blood samples were taken 45 min after a glucose injection, whereupon the level was seen to be significantly increased in the healthy mice (3.95±0.38 mmol/mg protein in glucose-injected mice versus 1.14±0.58mmol/mg protein in the placebo-injected mice) (Fig. 1c). Interestingly, a significantly high amount of glucose was detected in the skin of both the placebo and glucose-injected type I diabetic mice (27.35±3.5; and 25.67±4.3 mmol/mg protein; respectively) (Fig. 1c). To find the mechanism by which blood glucose is taken up by the skin, we examined the expression of glucose transporter-1 (Glut-1) at the mRNA and protein levels. Our data demonstrated a 26-fold increase in *Glut-1* mRNA in the healthy mice after the glucose injection (Fig. 1d), while the Glut-1 protein level was upregulated (Fig. 1e).

Examination of Glut-1 expression in the diabetic mice indicated that glucose injection affected neither its mRNA expression nor its protein expression (Fig. 1d, e). On the other hand, the *Glut-1* mRNA expression level in the placebo-injected type I diabetic mice was 3.4-fold higher than in the placebo-injected healthy ones (Supplementary Fig. 1a). This may explain why the glucose level is significantly higher in the skin of diabetic mice than in that of healthy mice. Taken together, these results suggest that a modification in blood glucose concentration will regulate the skin glucose level in the same direction.

### Trisk 95 is a novel biomarker for detecting glucose in the skin

To determine whether a specific pathway is activated in the skin upon an increase in blood glucose concentration, we performed label-free differential proteomic analyses of the skin biopsies taken from the placebo and glucose-injected healthy and type I diabetic mice. The results showed that Triadin isoform-1 (Trisk 95) was the only protein among all those screened (n=1824) that significantly increased in both the healthy and type I diabetic groups after glucose injection (Fig. 2a). To investigate further the effect of blood glucose concentration on Triadin expression, *Trisk 95* mRNA levels were measured in the skin of the different groups of mice.

**Fig. 2.**
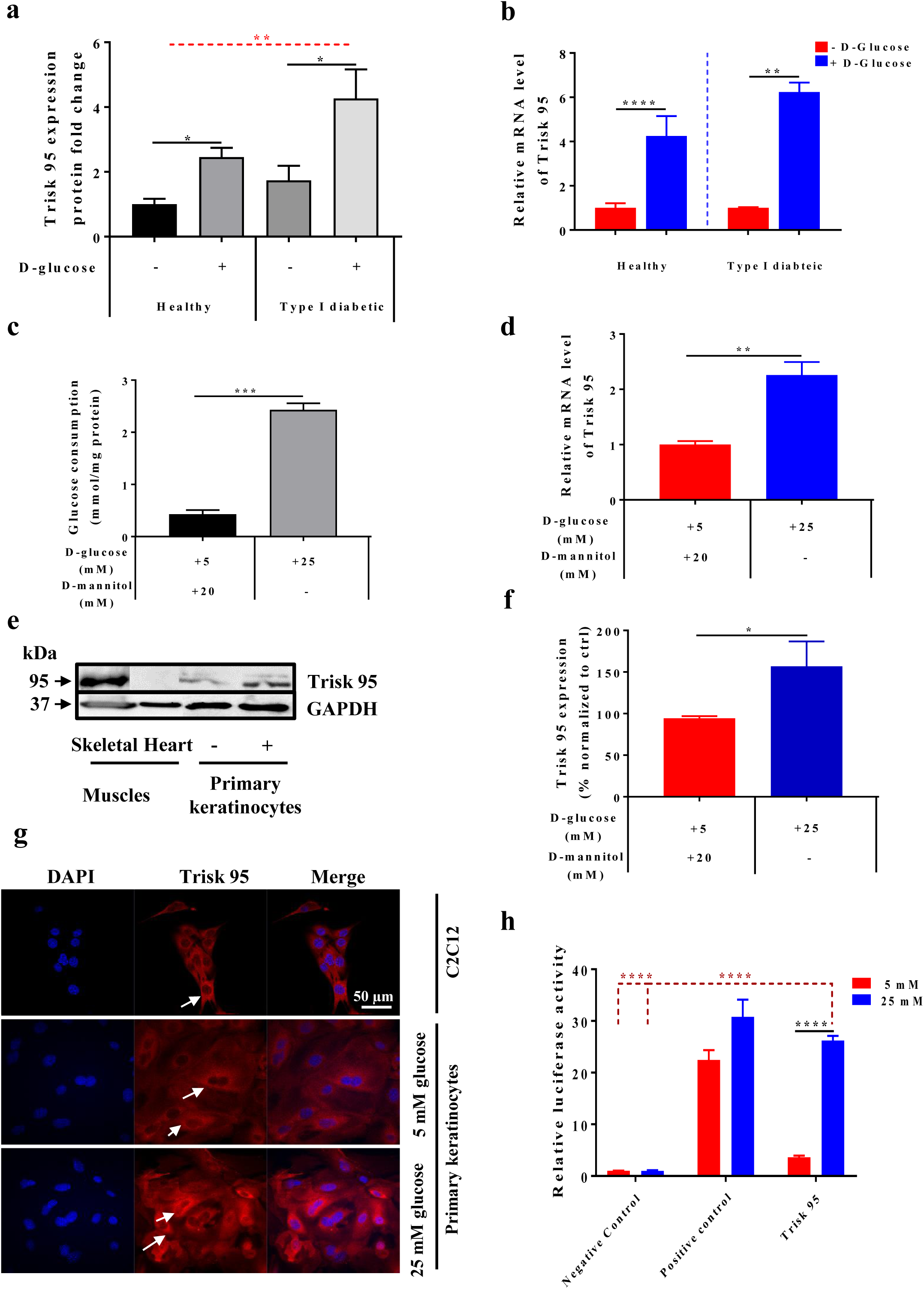
Trisk 95 is a new skin sensor for high blood glucose concentrations. **a** Proteomic profile of the skin after glucose injection *in vivo*. Skin biopsies from four groups were subjected to proteomic analysis. The results are presented as ratios of the fold change after normalization to the healthy mice injected with PBS (n=5/group). **b** Quantification of *Trisk 95* mRNA expression in the skin after high glucose using qRT-PCR. The results are shown as averages normalized to the control ± SD (n=6 in the healthy group and n=5 in the type I diabetic group). **c** Glucose consumption *in vitro* using primary keratinocytes. Glucose consumption was measured and normalized to protein concentrations. Data are presented as means ± SEM from three separate experiments. **d** Quantification of *Trisk 95* mRNA by qRT-PCR. The results are shown as averages after normalization to the control ± SEM in three separate experiments. **e** Western blot analysis of Trisk 95, positive (skeletal muscle) and negative (heart muscle) controls were used. A band at 95 kDa was detected in the positive control and primary keratinocytes, while no band was observed in the negative control. **f** Quantification of Trisk 95 protein expression levels in the primary keratinocytes. The results are shown as averages after normalization to the control ± SEM in three separate experiments. **g** Detection of Trisk 95 (red) and DAPI (blue) by immunofluorescence. The C2C12 cell line was used as a positive control. Trisk 95 expression was observed in the ER of myoblasts as well as in the keratinocytes (white arrows), the signal being stronger in the cells incubated with high glucose than in the controls. Scale bar = 50µm. **h** Triadin as a promotor of high glucose. Cells were transfected with the promotor construct and negative and positive control plasmids and incubated with low or high glucose for 48h. Relative luciferase activities are presented as means ± SEM after normalization to the negative controls (n=3 in separate experiments conducted in triplicate). One-way ANOVA was used for multiple comparisons in a (* *P* <0.02, ** *P* <0.002), the two tailed Student’s t-test was used in b (* *P* <0.02), in c (*** *P* <0.0002) and in d (* *P* <0.02). Two-way ANOVA multiple comparison was used in h (**** *P* <0.0001).

Consistent with the proteomic data, the results showed 3.45 to 4-fold increases in *Trisk 95* mRNA in both the healthy and type I diabetic mice receiving an intraperitoneal injection of glucose (Fig. 2b). Since the blood insulin concentration in type I diabetic mice does not change after a glucose injection of, we concluded that the upregulation of Trisk 95 in the skin mediated by an increase in blood glucose takes place via an insulin-independent pathway.

To further verify the effect of glucose on the Trisk level, we examined Trisk 95 expression in primary mouse keratinocytes cultured in the presence of low (5 mM) and high (25 mM) D-glucose concentrations. In this experiment, primary keratinocytes isolated from healthy mice were cultured in a low-glucose medium for 10 days. The medium was then replaced with a fresh medium containing either a low (5 mM) or a high (25 mM) glucose level. Forty-five minutes later cell and medium samples were collected for further analysis. Measurement of the glucose uptake indicated a significant increase in glucose consumption when the cells had been incubated for 45 min in the presence of a high glucose concentration (2.43±0.1mmol/ug protein versus 0.43±0.07mmol/ug protein) (Fig. 2c), while *Trisk 95* mRNA and protein expression levels had likewise increased 2.3 and 1.6-fold, respectively (Fig. 2d, e, f). Consistently with this, immunostaining of keratinocytes pointed to a significant increase in the expression of Trisk 95 (red) when cells were cultured in the presence of 25 mM D-glucose (Fig. 2g and Supplementary Fig. 2a, b).

To further examine whether the glucose concentration affected the transcriptional expression of Trisk, we used a luciferase reporter plasmid in which the 882 bp region upstream of the ATG translation initiation codon of human *TRDN* had been cloned upstream of the luciferase. The results showed a significant increase in luciferase expression upon treatment of the cells with 25 mM D-glucose (Fig. 2h), indicating that the extracellular glucose concentration affects Triadin expression at the transcriptional level.

### An increased blood glucose concentration will trigger intracellular calcium level modifications

Given that Trisk 95 is a well-known actor in calcium release from the endoplasmic reticulum (ER) in skeletal and heart muscles (31–34), we then wondered whether the Trisk 95 overexpression mediated by a high glucose concentration might affect the calcium level in the ER store. To answer this question, the basal level of intracellular calcium was imaged live using Fluo-4 AM 45 min after culturing primary mouse keratinocytes with either a low or a high-glucose medium.

To evaluate the release of calcium from ER, thapsigargin (an inhibitor of sacro/endoplasmic reticulum Ca2+ ATPase (SERCA) pump function) was added to block the pumping of calcium back into the ER. A significant increase in the maximum amplitude (background-subtracted F1/F0, where F0 is the minimum level of fluorescence before adding thapsigargin) was observed in the cells incubated with 25 mM D-glucose compared with the controls (0.32±0.037 versus 0.2±0.02) (Fig. 3a). We then measured the speed with which half of the maximum fluorescence amplitude was reached, and the results showed a significant difference between these two conditions, the speed being significantly slower in the cells incubated with high glucose (76.82±5.28 second) than in the controls (62.16±3.59 second) (Fig. 3b), suggesting that the release of calcium from the ER was higher under these conditions. Two representative movies showing the time lapse measurements with analysis of the regions of interest and the averaged fluorescence intensity traces under both sets of conditions are presented as supplementary data (Supplementary Fig. 3a and b).

**Fig. 3.**
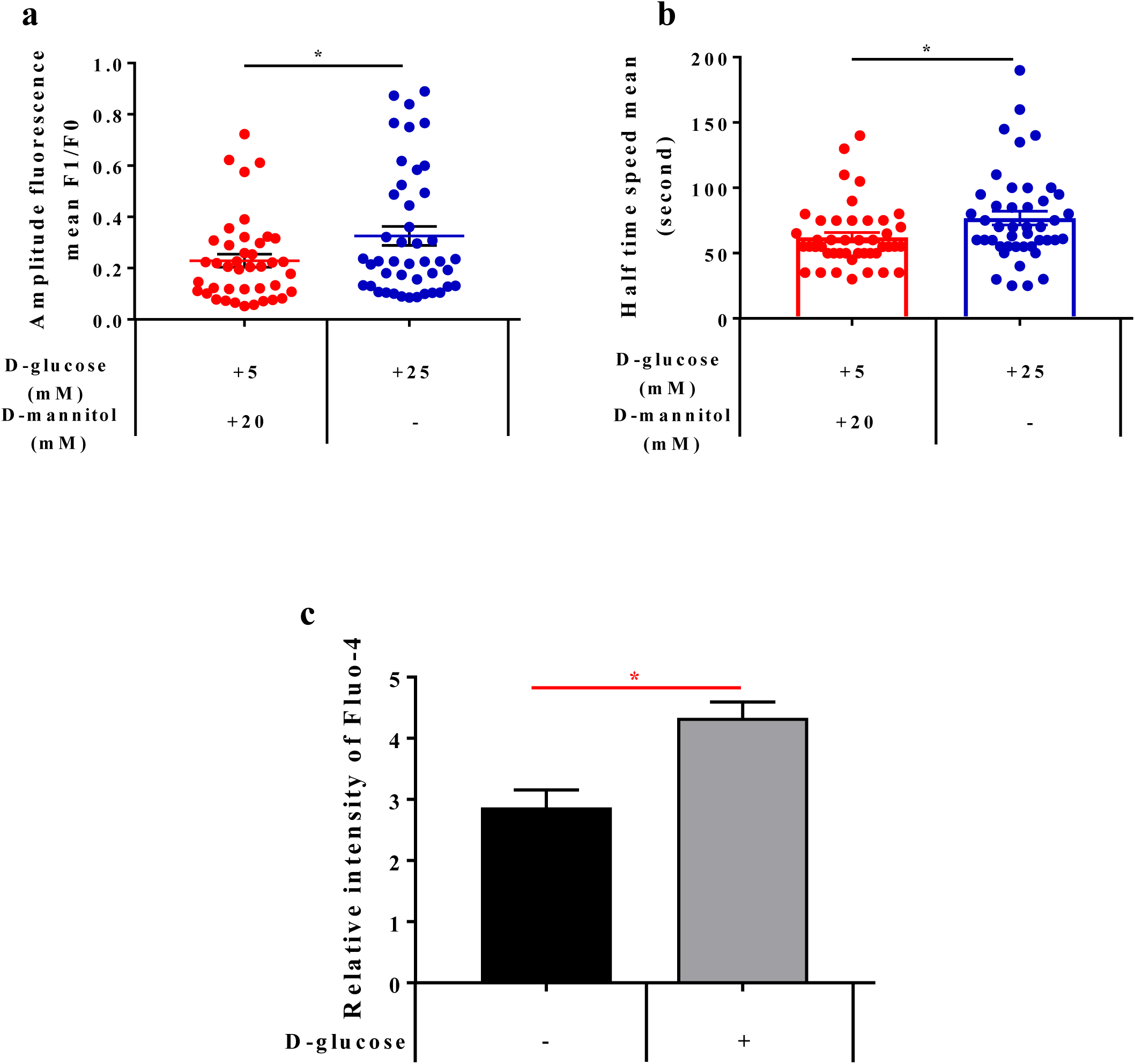
High extracellular glucose concentrations induce changes in the intracellular calcium in primary keratinocytes. **an** Intracellular calcium levels in primary keratinocytes after 45 min of glucose treatment. Cells were incubated for 30 min in the presence of Fluo-4 AM and intracellular calcium basal levels were measured for 50 seconds, after which thapsigargin was added to the medium. Time lapse recording was used to follow the increase in the fluorescence signal until the maximum amplitude was reached. The data showed a significant increase in the amount of cytosolic calcium in the cells incubated with high glucose relative to the controls (n=41). **b** Speed required to reach the half maximum of the fluorescence signal. The keratinocytes incubated with high glucose were slower to reach their peak than were the controls. **c** High extracellular glucose increased calcium uptake. The GTT test was performed using healthy mice. Keratinocytes were isolated from the skin and stained using Fluo-4 for 30 min. The relative intensity of staining was then measured by flow cytometry. The results are presented as means ± SEM (n=3 animals and 10000 cells/mouse). The two-tailed Student’s t-test was used for statistical analysis in a (* *P* <0.01), b (* *P* <0.02) and c (* *P* <0.019).

To test the *in vivo* effect of the blood glucose concentration on the calcium level in the skin, the GTT test was performed on the healthy mice. The relative basal calcium level in freshly isolated keratinocytes was then measured by flow cytometry using Fluo-4 AM. The results indicate that keratinocytes isolated from the glucose-injected mice had significantly higher basal calcium concentrations than cells taken from their control counterparts (4.33±0.26 versus 2.87±0.29 in the glucose and placebo-injected mice, respectively) (Fig. 3c).

### Primary keratinocytes exhibit alteration in mitochondrial network morphology due to the Trisk 95 upregulation

To test the hypothesis that mitochondria act to buffer intracellular calcium levels through the action of the mitochondrial calcium uniporter complex (34, 35), thereby affecting mitochondrial metabolism and the morphology of the mitochondrial network (36, 37), we first examined the effect of the extracellular glucose concentration on mitochondrial morphology using electron microscopy (EM). The results showed that the majority of the mitochondria were subjected to a modification in their morphology when cells were incubated with 25 mM D-glucose (Fig. 4a). For quantification purposes, we classified the mitochondria into two groups: 1 – a normal group that can take various shapes (elongated or round) with arranged cristae and an electron-dense matrix (yellow dotted circles) (Fig. 4a), and 2 – an abnormal group that are swollen, enlarged and round in shape, the disarrangement of their cristae accompanied by a partially or totally electron-lucent matrix (red dotted circles) (Fig. 4a).

**Fig. 4.**
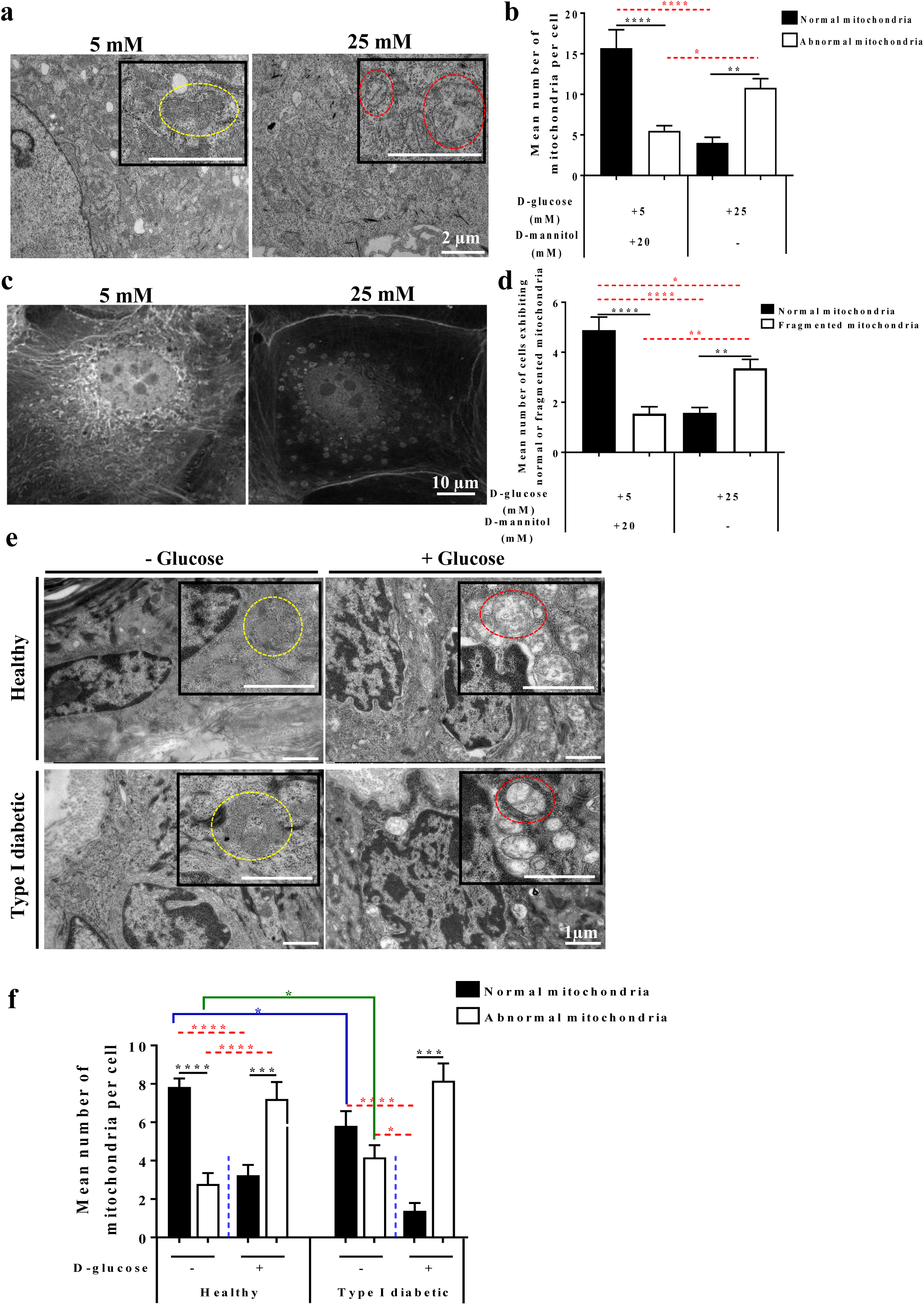
Keratinocytes exhibit a specific mitochondrial phenotype after high glucose treatment. **a** High extracellular glucose leads to an abnormal mitochondrial phenotype in keratinocytes. After 45 min of incubation in the presence of a low or a high-glucose medium, cells were fixed, and sections were examined using a Tecnai Spirit G2 transmission electron microscope. Keratinocytes incubated with high glucose demonstrated changes in the mitochondrial phenotype (swollen and enormous, round in shape, disarrangement of the cristae accompanied by partially or totally electron-lucent matrix) (red dotted circles); the normal mitochondria of different shapes (elongated or round) with well-arranged cristae and electron dense were observed in the cells incubated with low glucose (yellow dotted circles). Scale bar = 2µm. **b** Quantification of the mitochondria confirms the histological observations. A significant increase in the abnormal mitochondria was noted in the cells incubated with high glucose (n=45) compared with the controls (n=51). **c** The mitochondrial network in the keratinocytes was visualized using MitoTracker staining. A neat mitochondrial network surrounding the nucleus was observed in the control cells, but a fragmented network presented itself in the cells treated with high glucose. **d** Quantification of normal and fragmented mitochondria in control cells (n=328) and treated cells (n=316). **e** The morphology of the mitochondria was assessed in the skin after the GTT test. Histological examination of the mitochondria from the skin of healthy and type I diabetes mice demonstrated abnormal morphology (red dotted circles) after glucose injection similar to that observed in the primary cells, while the majority of the mitochondria of the control mice in the healthy and type I diabetes groups were healthy in shape (yellow dotted circles). Scale bar = 1µm. **f** Quantification of mitochondria in the skin. A significant increase in abnormal mitochondria was observed in the skin of both groups of healthy and type I diabetic mice injected with D-glucose (n=42 cells for all groups except n=35 cells for the type I diabetic mice injected with glucose) relative to their counterparts. One-way ANOVA multiple comparison was used for statistical analysis in b (* *P* < 0.03, ** *P* <0.004, **** *P* <0.0001), d (* *P <*0.015, ** *P <*0.001, **** *P <*0.0001 and f (**** *P* <0.0001).

Quantitative analyses indicated that while the majority of the mitochondria in the cells incubated with low glucose belonged to the normal group, the majority of those in the cells incubated with high glucose exhibited an abnormal shape (Fig. 4a, b). Since mitochondria always behave as a network (38, 39), we then assessed the effect of a high glucose concentration on the morphology of the mitochondrial network in keratinocyte mitochondria staining with the MitoTracker probe. While the majority of the control keratinocytes demonstrated an interconnected network located mostly around the nucleus, the majority of the cells incubated with high glucose exhibited a fragmented mitochondrial network that was dispersed in the cytoplasm (Fig. 4c, d).

To examine whether mitochondria are also affected under our *in vivo* conditions, skin biopsies were collected from the four groups of mice and visualized under EM. A similar mitochondrial phenotype was observed in the basal keratinocytes of both the healthy and type I diabetic groups upon receiving an injection of glucose. Indeed, while the majority of the mitochondria in the placebo-injected healthy group belonged into our normal mitochondrial class (yellow dotted circles) (Fig. 4e, f), the majority of those in both the glucose-injected healthy and type I diabetic groups were swollen, enlarged and round in shape with disarrangement of their cristae (red dotted circles) accompanied by a partially or totally electron-lucent matrix (Fig. 4e, f).

Significantly, the number of mitochondria belonging to the normal group was lower in the placebo-injected type I diabetic mice than in the placebo-injected healthy mice and that of the abnormal group higher (Fig. 4f). This could have been due to the high glucose level found in the skin of the type I diabetic mice even in the absence of glucose injection (see Fig. 1c).

### Trisk 95 knockout abolishes the increase in intracellular calcium mediated by a high glucose concentration and rescues the mitochondrial phenotype in the skin and its primary keratinocytes

To investigate the functional relation between Trisk 95 and intracellular calcium during the skin’s response to high blood glucose, skin biopsies were collected from the Trisk 95 KO mice 45 min after the injection of glucose or a placebo. To evaluate the basal calcium level, keratinocytes isolated from both groups were loaded with Fluo-4 AM. The results did not show any significant differences in basal calcium concentration between the placebo and glucose-injected Trisk 95 KO mice (2.74±0.62 versus 3.83±0.58, p=0.27) (Fig. 5a), indicating that Trisk 95 expression is necessary for high blood glucose-mediated modification to the skin calcium homeostasis.

**Fig. 5.**
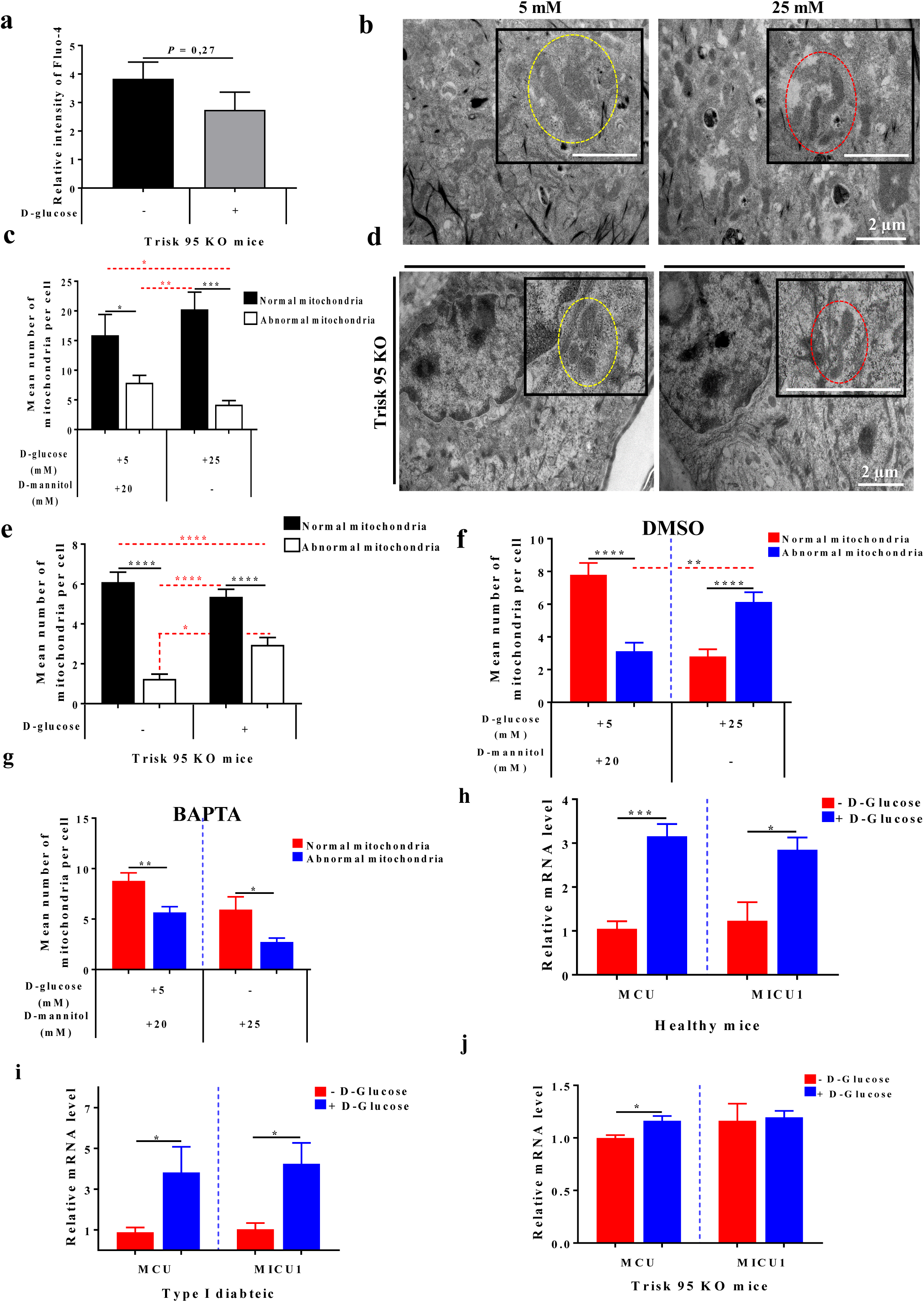
Trisk 95 KO rescues phenotype changes in both calcium and mitochondria in the skin and its primary cells. **a** Calcium uptake after high glucose injection was assessed. The relative intensity of Fluo-4 staining was measured by flow cytometry. The results are presented as means ± SEM (n=3 injected with PBS and n=4 injected with glucose and 10000 cells/mouse). **b and c** High glucose did not affect the mitochondria phenotype in Trisk 95 KO keratinocytes. Keratinocytes isolated from Trisk KO exhibited a normal mitochondrial phenotype (b), scale bar = 2µm. Quantification of mitochondria confirmed the histological observation (n=20) compared with the controls (n=29) (c). **d and e** High glucose did not affect the mitochondrial phenotype of Trisk 95 KO skin. The majority of the mitochondria from the Trisk 95 KO basal keratinocytes represented a normal phenotype (red dotted circles), as also observed in the controls (yellow dotted circles) (d), scale bar = 2µm. Quantification of the mitochondria in the skin injected with D-glucose (n=158 cells) as compared with the controls (n=132 cells) (e). **f** DMSO pre-incubation did not affect the mitochondrial phenotype brought about by high glucose. **g** BAPTA pre-incubation rescued the mitochondrial phenotype induced by high glucose in primary keratinocytes. **h, i and j.** Effect of high glucose on mitochondrial uniporter proteins in the skin. MCU and MICU1 expression in the skin. mRNA expression levels of *MCU* and *MICU1* in the skin of healthy (h), type I diabetic (i) and Trisk 95 KO (j) mice was examined using qRT-PCR. The results are shown as the averages after normalization to the controls ± SEM (n=6/group). The two-tailed Student’s t-test was used in a (ns. not significant *P* =0.27) and in e * *P* <0.01, ** *P* <0.009, *** *P* <0.0006. One-way ANOVA (multiple comparison) was used in d (* *P* <0.01, \*\*\*\**P* <0.0001), f (**** *P* <0.0001, *** *P* <0.0001, ** *P* <0.0014), and g (** *P* <0.003, * *P* <0.041) and the two-tailed Student’s t-test was used in h (*** *P* <0.0002, * *P* <0.012), i (* *P* <0.042 and * *P* <0.025) and j (* *P* <0.035).

In order to assess the role of Trisk 95 in the transitional modification of the mitochondrial phenotype, keratinocytes isolated from Trisk 95 KO mice and cultivated in the presence of low or high glucose were examined by EM. Interestingly, the majority of the mitochondria in the Trisk 95^-/-^ keratinocytes incubated with high glucose exhibited an elongated shape accompanied by arranged cristae and an electron-dense matrix (Fig. 5b, c), a phenotype that has been classified as normal. These results indicate that Trisk 95 expression plays an important role in keratinocyte responses to the extracellular glucose level. To examine this hypothesis further, skin mitochondrial phenotype comparisons were made between the placebo and glucose-injected Trisk 95^-/-^ mice, whereupon the results showed that the majority of the mitochondria in the basal keratinocytes were normal in both groups (Fig. 5d, e), indicating that Trisk 95 downregulation blocks the mitochondrial phenotype modification mediated by a high blood glucose concentration in.

To further examine the role of calcium in mitochondrial phenotype modification when monitoring high extracellular glucose concentrations, keratinocytes isolated from healthy mice were first incubated with BAPTA, a well-known calcium chelator, for 45 min and then subjected to low or high glucose conditions for 45 min. Histological examination of the mitochondrial phenotype indicated that treatment with DMSO (in which BAPTA was dissolved) did not affect the influence of extracellular glucose on the mitochondrial phenotype (Fig. 5f, see also Fig. 4b). The addition of BAPTA, however, significantly reduced the number of abnormal mitochondria in the cells subjected to high glucose (Fig. 5g). Taken together, our results indicate that the mitochondrial phenotype is triggered by calcium changes and can be rescued by adding BAPTA.

In order to analyse the molecular mechanism by which cytoplasmic calcium fluctuation could trigger mitochondrial phenotype alternation in the skin and its primary cells, we examined the expression of MCU and MICU1 (two well-known mitochondrial calcium uniporters) in the skin, and found that, upon subjection to high glucose injection, *MCU* and *MICU1* expression at the mRNA level were significantly increased in both healthy (3 and 2.8-fold increases, respectively) and type I diabetic mice (3.8 and 4-fold increases) compared with their control counterparts (Fig. 5h, i). Interestingly, when looking at MCU and MICU-1 expression levels in the Trisk 95 KO mice, we found that only *MCU* was significantly upregulated in the glucose-injected mice relative to the placebo-injected ones (a 1.16-fold increase) but not *MICU1* (Fig. 6j). These findings suggest firstly that MCU and MCU1 are required together for activation of the mitochondrial calcium uniporter complex in keratinocytes, allowing calcium uptake, and secondly that the main function of Trisk 95 in keratinocyte responses to high extracellular glucose is to orchestrate that activation.

**Fig. 6.**
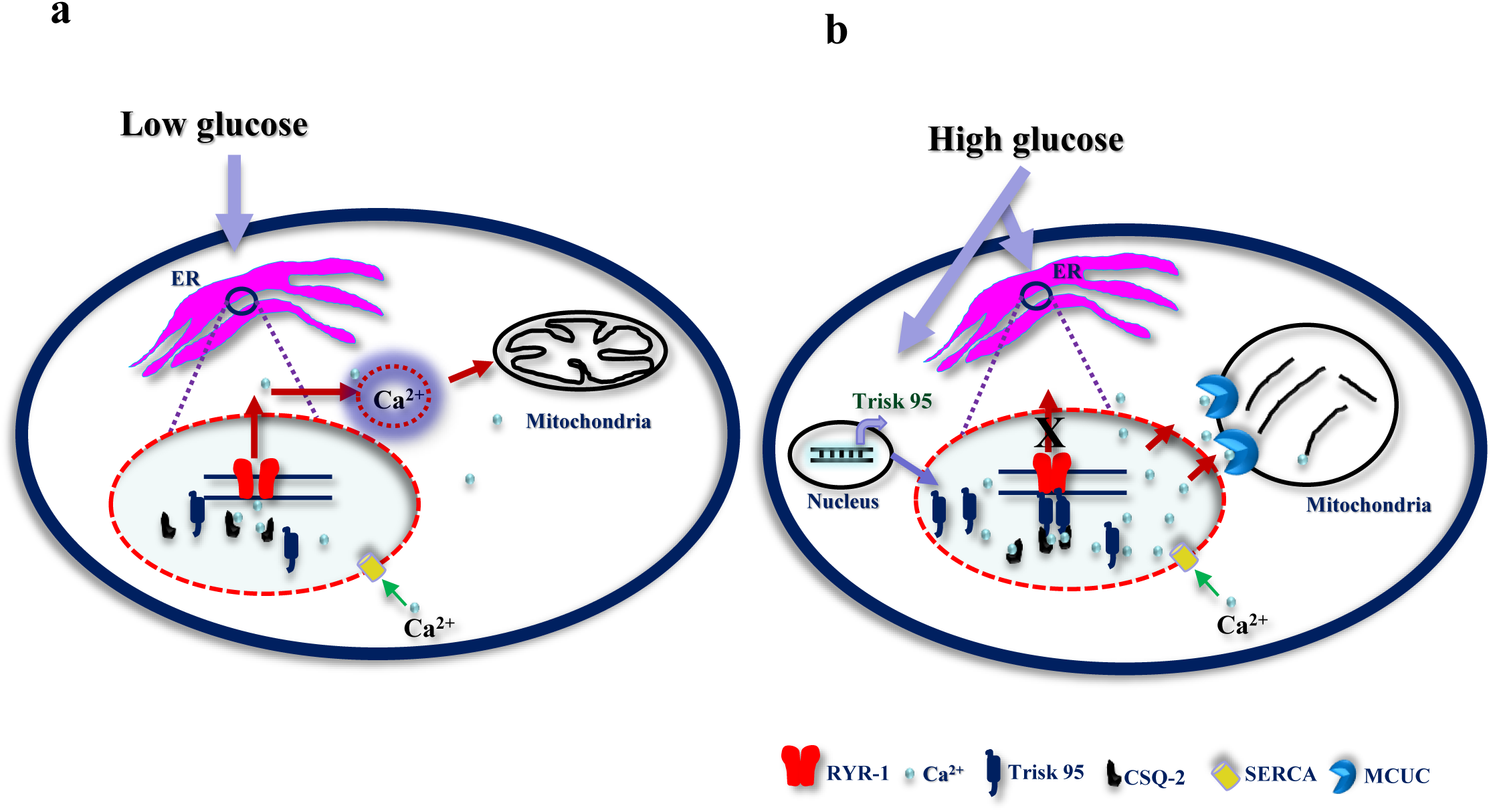
Proposed model for the mechanism by which high glucose triggers calcium uptake in the ER and mitochondria via Trisk 95. Under normal conditions (low glucose) no binding occurs between Trisk 95 and RYR and the calcium level in the ER is thus maintained via constant exchange with the cytosol. The calcium level is important for the functioning of the mitochondria (a). In the presence of high glucose Trisk 95 is overexpressed at transcriptional and then protein level, leading to its binding to the RYR channel and resulting in the closure of the latter. As a consequence, the increased calcium store in the ER triggers mitochondrial changes because of calcium uptake via MCUC (b).

## Discussion

We have shown here that the skin and its primary keratinocytes rapidly sense any increase in blood glucose concentration and trigger a response network.

Our results showed that upon an increase in blood glucose concentration the skin cells in both healthy and type I diabetic mice take up more glucose, suggesting that this takes place through an insulin-independent pathway. When looking for the mechanism underlying this increased uptake of glucose by keratinocytes we found that *Glut-1* mRNA expression was significantly increased in the mice receiving D-glucose compared with the controls, and that *Glut-1* mRNA expression was significantly higher in the type I diabetic mice than in the healthy mice, which may explain the higher skin glucose level.

A previous study has shown that keratinocytes were regulating their glucose uptake under hyperglycaemia by reducing the expression of the glucose transporter (Glut-1) (18). In that experiment, the keratinocytes were incubated for 5 days in a medium with either a low or a high glucose concentration, whereas in our protocol the period of incubation was 45 min. Our results indicate that keratinocytes are sensitive to the extracellular glucose concentration and that acute and chronic glucose sensing could lead to different responses. It is now important to understand how glucose is detected by keratinocytes and to know what the outcomes of increased glucose uptake by keratinocytes are.

Triadin is a key player in the functioning of the heart and skeletal muscles in that it regulates the excitation-contraction (EC) coupling process (32,40–42). Moreover, it modulates intracellular calcium haemostasis via its interaction with various proteins such as Junctin (43), RYR (44, 45) and CSQ (46). Indeed, it has been shown that Triadin regulates calcium release from the sarcoplasmic reticulum (SR) by binding directly to the RYR receptor in a specific domain (47). This interaction regulates RYR activity, leading to modification in the level of intracellular calcium in the myoblast cell line (48) and in the skeletal muscles cells (49). Our results show that a high extracellular glucose concentration induces Trisk 95 upregulation at the transcriptional level by regulating its promotor (Fig. 2h).

To test whether RYR and CSQ expression is modified under our conditions, we looked at their expression in our mice models. The results showed a significant increase in the *RYR-1* mRNA level in both glucose-injected healthy and type I diabetic mice relative to their counterparts (Supplementary Fig. 4a, b).

Regarding *CSQ-2* mRNA, its expression level was significantly increased in the glucose-injected type I diabetic mice (Supplementary Fig. 4a, b). Similar results were obtained from primary keratinocytes after 45 min of high glucose incubation (Supplementary Fig. 4c). Based on our results, we propose the following model for describing skin responses to fluctuations in blood glucose concentration (Fig. 6). In healthy individuals (blood glucose level ∼ 5 mmol/l), Trisk 95 does not bind RYR-1 or CSQ-2 and a constant calcium leak from ER into the cytosol is maintained via the RYR-1 channels. Under those conditions the cytoplasmic calcium is involved in physiological processes in the mitochondria (Fig. 6a). Under type I diabetic conditions, however, in which blood glucose levels are high (≥ 15 mmol/l), Trisk 95 is upregulated at both levels (mRNA and protein). Binding of the skin Trisk 95/CSQ-2 complex to the RYR-1 channels leads to their closure and subsequently to increased ER calcium stores. This increase in ER calcium eventually triggers modifications in the mitochondrial network and consequently in its optimal functioning (Fig. 6b).

Consistently with a critical role for Trisk 95 in orchestrating skin responses to an increase in the blood glucose concentration, increased calcium levels and modifications to the mitochondrial phenotype and network were absent in the glucose-injected Trisk 95 KO mice (Fig. 5a), and similarly no changes were observed when examining *RYR-1* and *CSQ-2* mRNA levels in Trisk 95 KO skin in response to a high-glucose injection (Supplementary Fig. 4d).

Our findings in these respects are in disagreement with previous data claiming a significant reduction in calcium release in Trisk 95 KO myotubes and muscles (32, 42). However, our data obtained from the overexpression of Triadin in keratinocytes confirm previous claims of higher resting SR calcium in heart muscle and myotubes cells (40, 50). In the light of our data and those of others, we speculate that Trisk 95 may be a key player in regulating the ER calcium store via regulation of the RYR channels.

It is worth mentioning that Inositol 1,4,5-trisphosphate receptor (IP3R) is playing role in regulating the calcium release from ER in keratinocytes (51–53). In addition, it has been reported that Triadin regulates IP3 receptors in rat skeletal myoblasts (54), further investigations to show whether possible interaction between Trisk 95 and IP3R might take place in keratinocytes.

It is known that a high concentration of calcium induces differentiation markers such as loricrin, profilaggrin and involucrin in keratinocytes (55–57), and it has also been shown that keratinocytes located in different layers of the epidermis express different RYR receptors (58). The effects of the calcium level and of RYR expression upon the upregulation of Triadin under hyperglycaemia may therefore explain the poor wound healing observed in type I diabetes.

Given that calcium signalling is a key regulator of mitochondrial behaviour (59), we hypothesize that an increase in intracellular and ER calcium may affect mitochondrial function. Our data obtained *in vivo* and *in vitro* indicate that mitochondria present a specific phenotype after high glucose in the skin of both healthy and type I diabetic mice. The mitochondrial phenotype might be attributable to calcium uptake resulting in mitochondrial damage (60). The mitochondrial calcium uniporter complex is responsible for calcium uptake across the inner membrane (34,35,61,62), and in agreement with this, our data showed that the expression of *MCU* and *MICU1* at the mRNA level was significantly upregulated after high glucose in both healthy and diabetic mice (Fig. 5h, i).

Relationships between calcium and mitochondrial morphology changes have been reported in astrocytes (63, 64) and in human keratinocytes (HaCat) (65), but interestingly; data published by Deheshi S *et al*., demonstrated that calcium-induced ROS production could plausibly be the mechanism underlying the mitochondrial morphological changes observed in astrocytes (elongated to uniformly punctate organelles) and their fragmentation (64).

The effect of a high glucose concentration on mitochondrial fragmentation has been reported in several cell types, such as retinal endothelial cells (66), a liver cell line (67, 68) and neonatal rat ventricular myocytes (69). *Trudeau et al*. have shown that an increased glucose concentration will induce bi-phasic changes in mitochondrial morphology in retinal pericytes (70). Indeed, a transient modification has been seen to occur during the first 6h, followed by a second modification starting from 48h and lasting until 7 days in the presence of high glucose (70). In view of the literature and our present results, we speculate that increased calcium may induce changes in ROS production in our model, so that further investigations should be considered.

Overall, our study highlights the rapid, dynamic physiological behaviour occurring in the skin and its primary keratinocytes in response to blood glucose concentration. The data demonstrate for the first time that high blood glucose for 45 min induces the activation of a number of players that are cross-talking, leading to the activation of signalling cascades. Trisk 95 is shown to be an essential mediator in the interaction between these cascades and proposes that the ensuing calcium can be used as an indicator of the blood glucose level in the skin. This work represents the first step of our registered patent (No. PCT FI2016/050917, https://encrypted.google.com/patents/WO2017109292A1?cl=no), and suggests further studies for developing the device that is able to detect the biological changes in the cells and convert them to a value.

## Materials and Methods

### Cell culture and glucose treatment

Primary keratinocytes were isolated from the skin of 6-8-week-old mice as previously described (Supplementary ref.^1^). Briefly, mice (C57/BL6) were sacrificed (by cervical dislocation) and the hair shaved from their backs. The skin was then sterilized by immersion in a beaker of 10% betadine for 2 min, in a beaker of 70% ethanol for 1 min and then in sterile Dulbecco’s phosphate (PBS) for 1 min. The skin samples were treated with trypsin for 2h at 33°C to separate the epidermis from the dermis. Keratinocytes were cultured in F12 (21765-037) – DMEM (41965-062), medium, both from Thermo Fisher, supplemented with hydrocortisone (0.5 µg/ml), epidermal growth factor (EGF) and insulin (5 µg/ml), all from Sigma. The medium was changed three times a week. When the cultures reached 70– 80% confluence the keratinocytes were incubated with two concentrations of D-glucose (G7021-Sigma) (5 mM + 20 mM mannitol (for osmolality)) or 25 mM for 45 min and then used for the various experiments.

### The type I diabetic mouse model

Type I diabetes was induced after one single intraperitoneal (IP) injection of streptozotocin (STZ), as previously described (30). One week before the experiment, all the mice were housed in individual cages. On the day of the experiment, they fasted for 4h and were then divided into two groups: controls and type I diabetic mice. The control mice were injected IP with Na citrate buffer (pH 4.5), while the type I diabetic mice were injected with STZ (150 mg/kg) diluted in Na citrate buffer to a total volume of 100µl. The mice were diagnosed as diabetic when their blood glucose level was ≥ 15 mmol/l.

### Insulin measurement

To determine the insulin concentration in the serum, blood samples were collected from the facial vein of the mice 45 min after the D-glucose injection. Serum was obtained after centrifugation at 3500 rpm for 10 minutes at 4° C and insulin was measured using the Insulin Rodent (Mouse/Rat) Chemiluminescence ELISA kit (ALPCO) according to the manufacturer’s instructions

### Glucose tolerance test

For adaptation purposes, the mice were housed in individual cages for one week. Prior to the experiment the healthy and type I diabetic mice fasted overnight (12h) and each group was then divided into two parts, yielding four groups in all: healthy and type I diabetic groups that were injected intraperitoneally (IP) with PBS, and healthy and type I diabetic groups that were injected IP with D-glucose (2g/kg). Blood glucose levels were monitored at 0, 5, 15, 30- and 45-min post-injection. After 45 min, all the mice were killed and skin samples were collected for the various experiments (RT-qPCR, western blotting and EM). All the animal procedures were performed in accordance with the European Convention (ETS 123), EU Directive 86/609/EEC and the Finnish national legislation, as approved by the local ethical committee.

### RNA extraction from skin samples and primary keratinocytes

Total RNA was extracted from the skin using Trizol (Invitrogen, 15596-026) and subsequently the RNeasy Mini Kit (Quiagen). The tissues were cut into small pieces, incubated with Trizol for 5 min at 4°C and homogenized using TissuLyser^TM^ (Qiagen) for 5 min at 50 Hz. Skin lysates were centrifuged for 15 min at 13000 rpm at 4°C, after which chloroform was added to the supernatant and the samples incubated for 5 min at RT followed by centrifugation for 18 min at 13000 rpm at 4°C. The rest of the extraction steps were performed using the RNeasy Mini Kit according to the manufacturer’s instructions. The RNeasy Mini Kit was also used to purify the RNA from the primary keratinocytes. The quality of the RNA from both the skin and the cells was assessed using nanodrop.

To analyse the mRNA expression level of *Trisk 95*, 2 μg of RNA was reverse transcribed to cDNA using the SuperScript™ VILO™ cDNA Synthesis Kit (ThermoFisher, 11754050) or the First Strand cDNA synthesis kit (Roche Applied Science). Quantitative real-time PCR was carried out for *Glut-1*, *Trisk 95*, *RYR-1*, *CSQ-2*, *MCU*, *MICU1*, *18S* and *GAPDH* using primers from a TaqMan gene expression assay and the SYBER Green method. The primer sequences used are listed in Table 1. The reactions were cycled 40 times after initial polymerase activation (50°C, 2 minutes) and initial denaturation (95°C, 20 minutes) using the following parameters: denaturation at 95°C for 1 second, and annealing and extension at 60°C for 20 seconds. The relative expression of target genes was normalized to *18S* or *GAPDH* expression and the ΔΔCT method was applied followed by a two-tailed Student’s t-test.

**Table 1.**
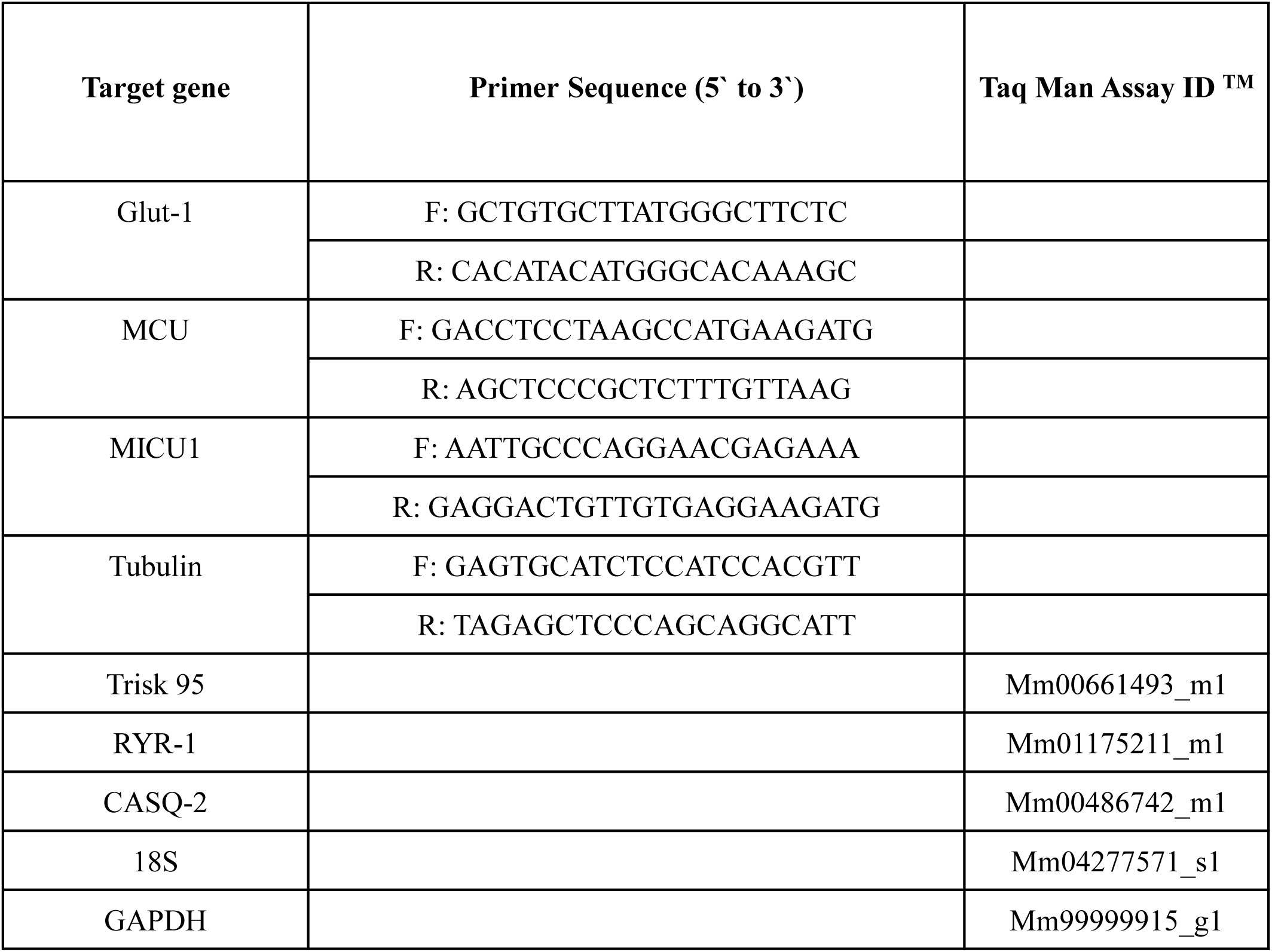
Primer sequences for quantitative real-time PCR.

### Luciferase reporter assay

The *Triadin* promoter region containing a 882 bp (chr6:123636957+123637838) DNA fragment was cloned into the pGL3 basic vector (Promega) upstream of the luciferase reporter gene. HaCat cells were seeded in a 96-well white plate at a concentration of 2×10^4^ in the presence of 100 μl DMEM (41965039 gibco) medium supplemented with 10% FBS and 1% PS. 12 hours later, the medium was replaced with 100 µl of DMEM-free serum supplemented with 1% PS containing 5mM D-glucose + 20mM D-mannitol or 25mM D-glucose (Sigma). The cells were then immediately transfected with a promoter construct, negative and positive control plasmids (100ng each) and an internal control plasmid pGL4.74 (40ng) (Promega), using Turbofect (0533 Thermofischer Scientific) as the transfection reagent according to the manufactureŕs protocol. The cells were incubated at 37℃ in 5% CO. Two days later the Dual-Glo Luciferase Assay System (E2940, Promega) was used to measure the luciferase activity of the cells on a bio-luminometer, following the manufactureŕs protocol.

To avoid interference from firefly luminescence, all the samples on the 96-well plate were measured before inducing renilla luminescence. All the readings were based on measurements obtained from at least three replicate wells. The change in the activity of each gene promotor was determined by measuring the activity of luciferase in the transduced keratinocytes incubated with high glucose (25 mM) versus those incubated with low glucose (5 mM). Statistical significance was analysed using the two-tailed Student’s t test. Each experiment was performed independently three times. The following primers were used for cloning the Triadin promoter region (882 bp):

Triadin-882-NheI-F CTAGCTAGCTTCTCCCCCTGGAAACAACC

Triadin-882-XhoI-R GTACTCGAGCACCTGTCTAGGAGTGG

### Proteomic analysis

Details of the sample preparation, nano-scale liquid chromatographic tandem mass spectrometry analysis, database search, processing of the results and label-free quantitative data analysis are provided in the Supplementary materials section.

### Western blotting

Equal amounts of total protein were resolved by SDS-PAGE and electrophoretically transferred to nitrocellulose membranes (BioTop 741280). The membranes were then incubated overnight at 4°C with a 1:1000 dilution of anti-Trisk 95 (Supplementary ref.^2^), Glut-1 (Supplementary ref. ^3^) 1:5000 anti-GAPDH (MAB374, Millipore), 1:5000 anti-β-actin (A5441, Sigma Aldrich). After treatment with peroxidase-linked antibody (Dako, polyclonal) for 1 hour, blots were developed using the 20X LumiGLO® Reagent and 20X Peroxide (Cell Signaling Technology, #7003).

### Measurement of glucose uptake and intracellular glucose consumption

To evaluate glucose uptake or its intracellular concentration, we used the YSI 2950 Biochemistry Analyzer (YSI Life Sciences), which employs immobilized enzyme to catalyse the corresponding chemical reactions in order to measure the concentration of a specific metabolite. Briefly, to measure glucose uptake, keratinocytes were incubated in triplicate for 45 min in a 24-well plate in 1 ml of DMEM complete medium supplemented with glucose (5 mM or 25 mM). The concentration of glucose in the collected medium was evaluated by the YSI analyser using corresponding fresh medium as a reference.

To measure intracellular glucose concentrations, skin samples taken from the various groups of mice were immersed in a hypotonic buffer (2.5 mM Tris/HCl, pH 7.5 and 2.5 mM MgCl2) on ice for 15 min, homogenized and then sonicated for 15 s. The concentrations of glucose in the skin lysates were then evaluated. The data were expressed as mmol·μg−1 of protein.

### Intracellular calcium imaging and measurement

To image and measure intracellular calcium changes *in vivo*, primary keratinocytes were isolated and cultured in dishes with a glass bottom (35/10 MM, Greiner Bio-one International, 627871) for one week. On the day of the experiment the cells were washed and incubated for 45 min in the presence of 5 mM D-glucose + 20 mM mannitol or 25 mM D-glucose. Then Fluo-4 AM (2 µM final concentration) (Thermo Fisher, F14201) was added directly to the medium and incubated for another 30 min at 37℃. Changes in the intracellular calcium level was calculated by measuring the background-subtracted fluorescence signals (F1) and dividing these by the baseline signal (F0) before adding thapsigargin (Sigma, T9033) to a final concentration 100 nM. Zeiss Cell Observer Microscope (Carl Zeiss, Germany) in epifluorescence imaging mode was used to detect and measure the fluorescence of Fluo-4. The data were analysed and calculated by using Zen 2012 (Blue), the 2018 version of Origin and presented as means ± standard deviation.

To assess calcium levels after GTT, keratinocytes were isolated from skin biopsies collected at 45 min and stained Fluo-4 AM was added to the medium and incubated in the dark for 30 min. After two washes with PBS, the cells were analysed by flow cytometry. Data from ten thousand individual cells were collected for each sample.

### Immunostaining of cells

To examine the expression of Trisk 95 and its localization in the primary keratinocytes, cells were cultured in dishes with a glass bottom for 45 min after treatment with two concentrations of glucose. They were then fixed in 4% paraformaldehyde (PFA) for 10 min at RT and incubated with blocking solution (10% BSA, TBST (Tris buffer saline-Tween) 0.1%) for 30 min. This was followed by a further incubation with the primary antibody anti-Trisk 95, as previously described (Supplementary ref.^2^), diluted in the blocking solution for 1h at RT and then with DAPI, and with a secondary antibody for 30 min. Fluorescence was detected using a confocal microscope (Zeiss LSM 780, Carl Zeiss, Germany).

### Transmission electron microscopy (TEM)

Skin biopsies were collected from the four groups (three mice per group) after the GTT assay, and cultured cells were mounted on coverslips and fixed in 1% glutaraldehyde and 4% formaldehyde in 0.1 M phosphate buffer, pH 7.4. The remaining steps were followed as previously described (Supplementary ref.^4^). Sections from the skin samples and cells were examined using a Tecnai Spirit G2 transmission electron microscope. Images were captured with a Veleta CCD camera and Item software (Olympus Soft Imaging Solutions GMBH, Munster, Germany).

### Statistical analysis

GraphPad Prism software, version 7, was used for the statistical analyses. The one-way or two-way ANOVA test or the two-tailed Student t-test were employed and \**P*-values less than 0.05 were considered significant.

## Acknowledgements

We thank Dr. D. Vicente for his help in performing the GTT and Dr. G. Bart for her assistance and helping in extracting RNA from the skin tissues. We thank Dr. I. Skovorodkin for his scientific advice, J. Kekolahti-Liias and P, Haipus for their excellent technical assistance in the laboratory, and S. Tausta for her expert assistance and advice concerning GTT and the type I diabetes mouse model. We thank Dr. I. Miinalainen for his help with imaging the mitochondria using electron microscope. We are also grateful to Dr. M. Kaakinen for the C2C12 cell line and to Prof. K. Hiltunen and Docent A. Kastaniotis for their help with the mitochondrial phenotype. This work was supported Academy of Finland grants (251314; 315030; 307533; 206038; 121647), Finnish Cultural, Sigrid Jusélius and Finnish Cancer Research Foundations, the European Community’s Seventh Framework Programme Health (FP7/2007-2013) under grant agreement FP7-HEALTH-F5-2012-INNOVATION-1 EURenOmics 305608, and H2020-FETOPEN-2018-2019-2020-01 projects MindGap GA 829040 and Gladiator (GA 828837).

Supplementary Figures

**Supplementary Figure 1.**
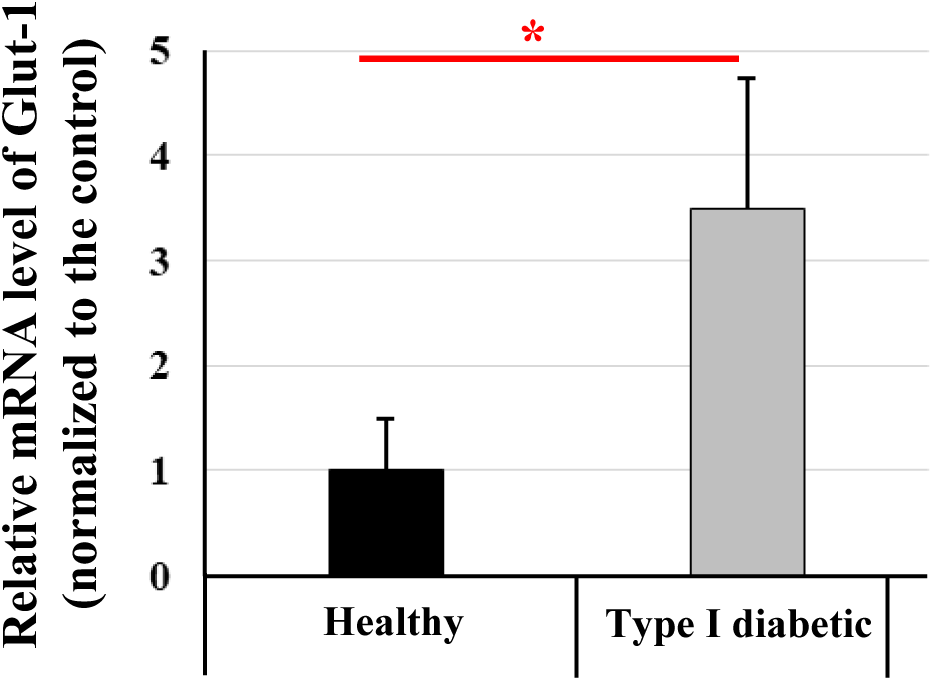
Quantification of the *Glut-1* mRNA expression in the skin of type I diabetic and healthy mice receiving PBS injections. The results are presented as average percentages after normalization to the controls ± SD. n=4/group, * *P* <0.03

**Supplementary Figure 2.**
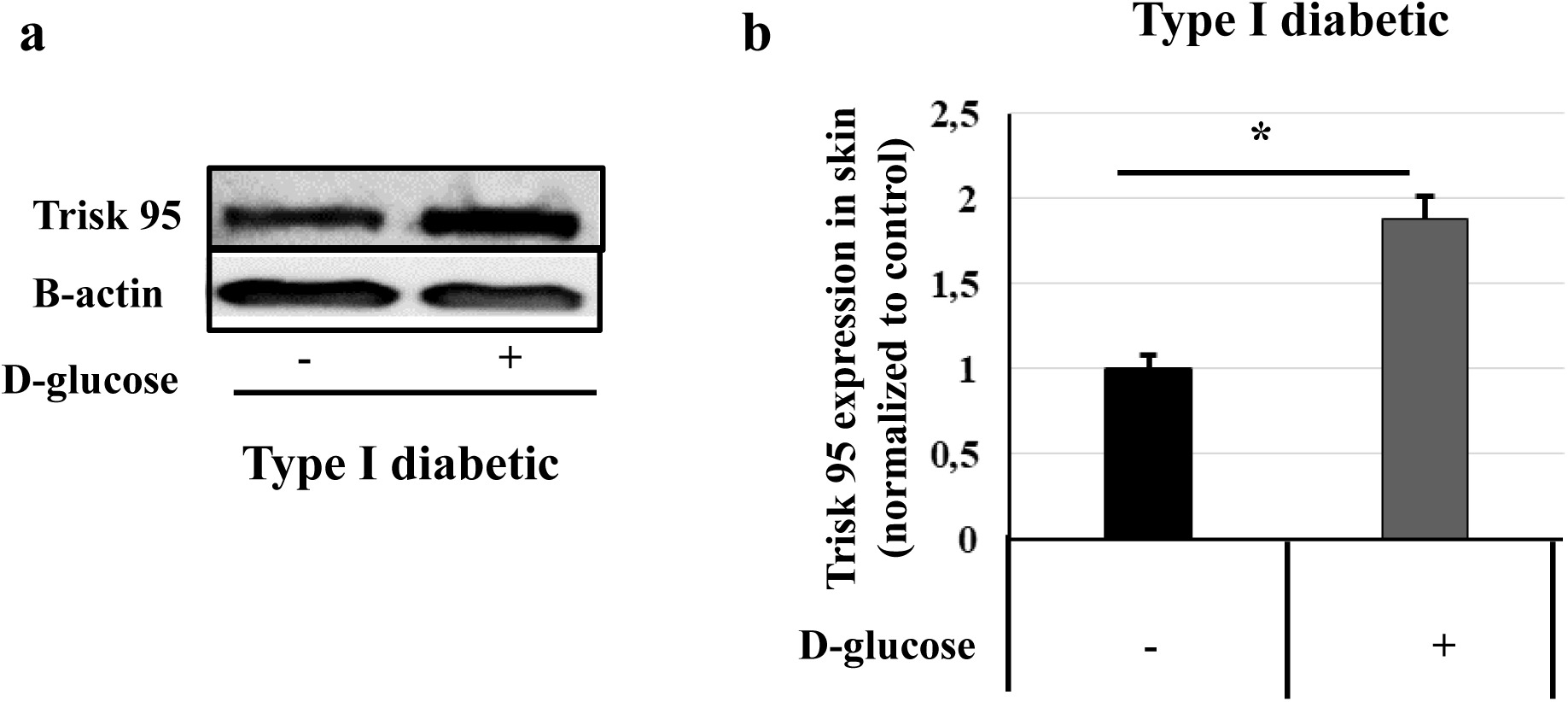
Triadin expression in skin from type I diabetic mice. **a** Western blot analysis of Trisk 95 in the skin, showing a band at 95 kDa. **b** Quantification of the Trisk 95 band in the skin of type I diabetic mice injected with high glucose. The results are presented as average percentages after normalization to the controls ± SD. n=4 and 5, * *P* <0.03.

**Supplementary Figure 3.**
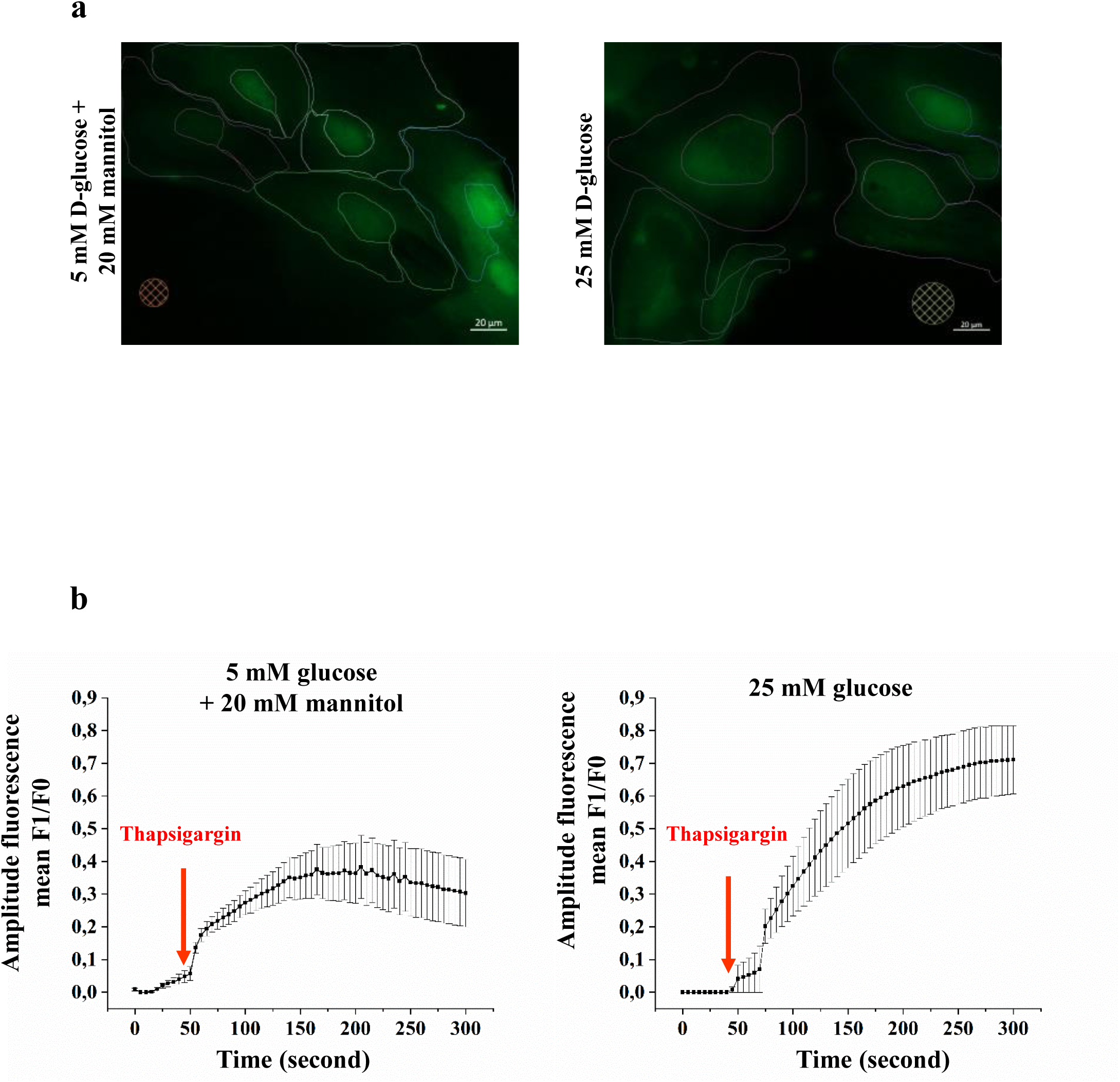
Calcium release in primary keratinocytes. Cells were incubated with low or high glucose for 45 min, and then stained with Fluo-4 AM for 30 min. Live-cell time lapse imaging was used to evaluate calcium release from the ER after blocking SERCA with thapsigargin. **Two movies of keratinocytes** incubated with 5 mM D-glucose and 25 mM D-glucose, showing the Fluo-4 intensity changes after adding thapsigargin at 50 seconds. **b** Averaged fluorescence intensity curves in the cells incubated with low or high glucose before and after thapsigargin stimulation. The cytosolic fluorescence intensity was measured within the regions of interest excluding the nuclei in each cell. The maximum fluorescence amplitude and the increase in speed (of reaching half maximum amplitude) was measured under both sets of conditions.

**Supplementary Figure 4.**
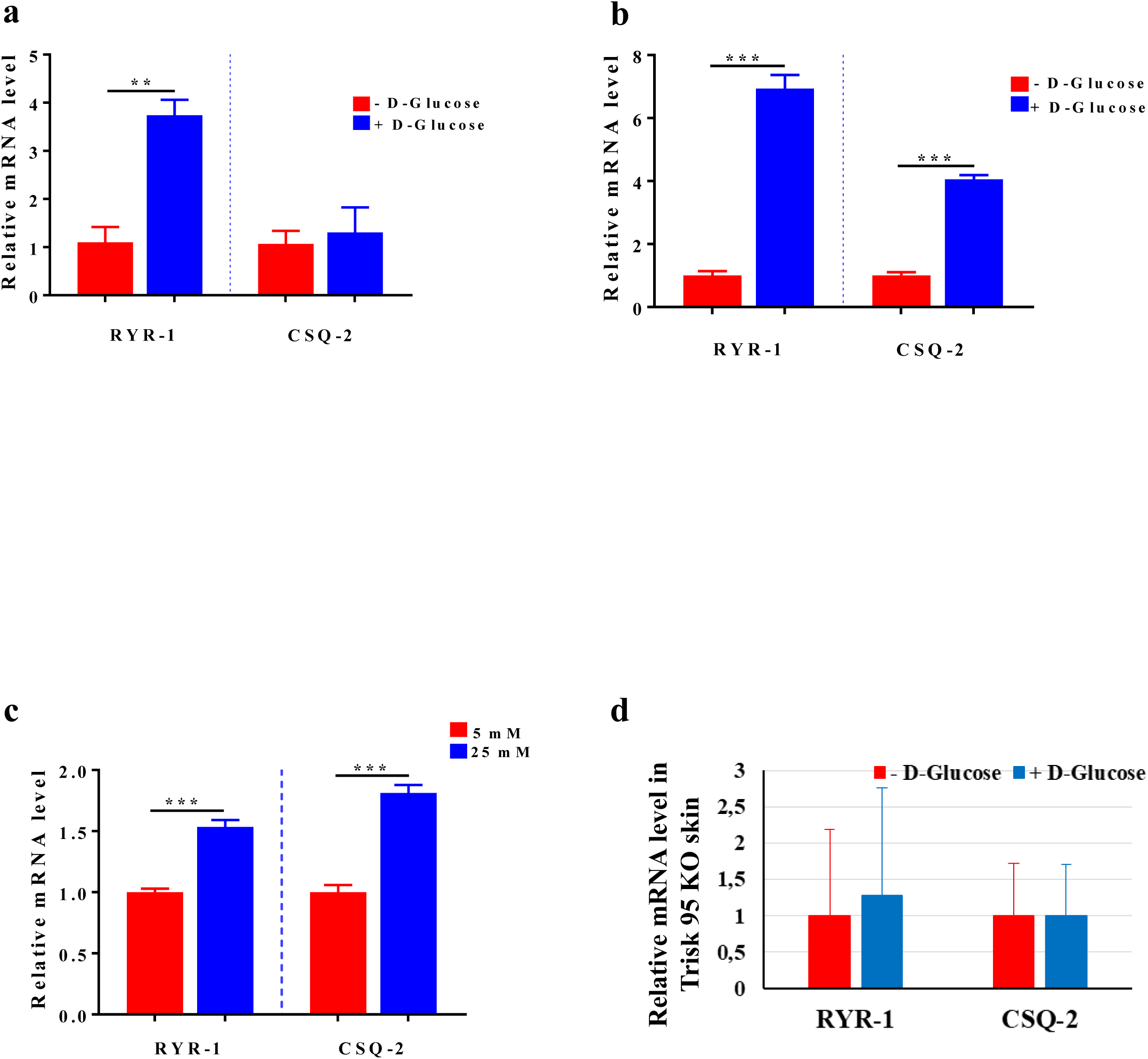
High glucose induces changes in RYR-1 and CSQ-2 expression, while Trisk 95 silencing will rescue their expression. **a, b and c** RYR-1 and CSQ-2 expression in the skin and primary keratinocytes. mRNA expression levels of *RYR-1* and *CSQ-2* in healthy mice (a), type I diabetic mice (b) and primary keratinocytes (c). **d** RYR-1 and CSQ-2 expression in Trisk 95 KO mice. The results are shown as averages after normalization to the controls ± SEM (n=6/group of healthy mice, n=3/group of type I diabetic mice, and three independent experiments for the primary keratinocytes. Also, n=3 (Trisk 95 KO mice injected with PBS) and n=4 (Trisk 95 KO mice injected with glucose). The two-tailed Student’s t-test was used for statistical analysis. ** *P* <0.004, n.s *P* =0.70 (a), *** *P* <0.001 (b) *** *P* <0.001 (c) and n.s, *P* =0.2 (d).

## Supplementary materials and methods

### Proteomic analysis

#### Sample preparation and protein digestion

Ten µg of each protein sample were solubilized in Laemlli buffer and deposited on SDS-PAGE gel for concentration and cleaning purposes. Separation was stopped once the proteins had entered the resolving gel. After colloidal blue staining, bands were cut out from the SDS-PAGE gel and subsequently cut into 1 mm x 1 mm pieces. These gel pieces were destained in 25 mM ammonium bicarbonate, 50% ACN, rinsed twice in ultrapure water and shrunk in ACN for 10 min. After ACN removal, the gel pieces were dried at room temperature, covered with trypsin solution (10 ng/µl in 40 mM NH4HCO3 and 10% ACN), rehydrated at 4°C for 10 min, and finally incubated overnight at 37°C. Spots were then incubated for 15 min in 40 mM NH4HCO3 and 10% ACN at room temperature with rotary shaking.

The supernatant was collected, and an H2O/ACN/HCOOH (47.5:47.5:5) extraction solution was added to the gel slices for 15 min. The extraction step was repeated twice, and the supernatants were pooled and concentrated in a vacuum centrifuge to a final volume of 100 µL. The digests were finally acidified by the addition of 2.4 µL of formic acid (5%, v/v) and stored at −20 °C.

### nLC-MS/MS analysis

Peptide mixtures were analysed on an Ultimate 3000 nanoLC system (Dionex, Amsterdam, Netherlands) coupled to an Electrospray Q-Exactive Quadrupole Orbitrap benchtop mass spectrometer (Thermo Fisher Scientific, San Jose, CA). Ten microlitres of peptide digests were loaded onto a 300-µm inner diameter x 5-mm C_18_ PepMapTM trap column (LC Packings) at a flow rate of 30 µL/min. The peptides were eluted from the trap column onto an analytical 75-mm id x 25-cm C18 Pep-Map column (LC Packings) with a 4–40% linear gradient of solvent B in 108 min (solvent A was 0.1% formic acid in 5% ACN, and solvent B was 0.1% formic acid in 80% ACN). The separation flow rate was set at 300 nL/min.

The mass spectrometer operated in positive ion mode at a 1.8-kV needle voltage. Data were acquired using Xcalibur 2.2 software in a data-dependent mode. MS scans (*m/z* 350-1600) were recorded at a resolution of R = 70 000 (@ m/z 200) and an AGC target of 3 x 10^6^ ions collected within 100 ms. Dynamic exclusion was set to 30 s and the top 12 ions were selected after fragmentation in HCD mode. MS/MS scans with a target value of 1 x 10^5^ ions were collected with a maximum fill time of 100 ms and a resolution of R = 17500. Only +2 and +3 charged ions were selected for fragmentation. The other settings were as follows: no sheath nor auxiliary gas flow, heated capillary temperature 250°C, normalized HCD collision energy 25% and isolation width 2 m/z.

### Database search and processing of the results

Data were searched by SEQUEST through Proteome Discoverer 1.4 (Thermo Fisher Scientific Inc.) against a subset of the 2016.07 version of the UniProt database restricted to the Mus musculus Reference Proteome Set (49153 entries). Spectra from peptides higher than 5000 Da or lower than 350 Da were rejected. The search parameters were as follows: mass accuracies of the monoisotopic peptide precursor and peptide fragments were set to 10 ppm and 0.02 Da, respectively.

Only b- and y-ions were considered for the mass calculations. The oxidation of methionines (+16 Da) was considered to be a variable modification and the carbamidomethylation of cysteines (+57 Da) to be a fixed modification. Two missed trypsin cleavages were allowed. Peptide validation was performed using the Percolator algorithm (Supplementary ref.^5^) and only “high confidence” peptides were retained, corresponding to a 1% False Positive Rate at the peptide level.

### Label-Free Quantitative Data Analysis

Raw LC-MS/MS data were imported in Progenesis QI for Proteomics 2.0 (Nonlinear Dynamics Ltd, Newcastle, U.K). Data processing includes the following steps: (i) Features detection, (ii) Features alignment across 12 samples, (iii) Volume integration for 2-6 charge-state ions, (iv) Normalization on total protein abundance, (v) Import of sequence information, (vi) Calculation of protein abundance (sum of the volume of corresponding peptides), (vii) A t-test was calculated for each group comparison and proteins were filtered based on a p-value<0.05. Significantly, only non-conflicting features and unique peptides were considered for calculation at the protein level. Quantitative data were considered for proteins quantified in a minimum of 2 peptides.

ACN Acetonitrile

Da Dalton

CID Collision Induced Dissociation

SDS Sodium dodecyl sulphate

